# Putative novel avian paramyxoviruses identified from wild bird surveillance (United States, 2016‒2018) and retrospective analysis of APMV-2s and APMV-6s in GenBank

**DOI:** 10.1101/2021.11.12.468463

**Authors:** Kelsey T. Young, Jazz Q. Stephens, Rebecca L. Poulson, David E. Stallknecht, Kiril M. Dimitrov, Salman L. Butt, James B. Stanton

## Abstract

Avian paramyxoviruses (APMVs) (subfamily *Avulavirinae*) have been isolated from over 200 species of wild and domestic birds from around the world. The International Committee on Taxonomy of Viruses (ICTV) currently defines 22 different APMV species, with *Avian orthoavulavirus 1* (whose viruses are designated as APMV-1) being the most frequently studied due to its economic burden to the poultry industry. Less is known about other APMV species, including limited knowledge on the genetic diversity in wild birds and there is a paucity of public whole genome sequences for APMV-2 to -22. The goal of this study was to use MinION sequencing to genetically characterize APMVs isolated from wild bird swab samples collected during 2016–2018 in the United States. Multiplexed MinION libraries were prepared using a random strand-switching approach using 37 egg-cultured, influenza-negative, hemagglutination-positive samples. Thirty-five APMV isolates that had complete polymerase coding sequences were speciated using ICTV’s current *Paramyxoviridae* phylogenetic methodology. Viruses from APMV-1, -4, -6, -8 were classified, one putative novel species (*Avian orthoavulavirus 23*) was identified from viruses isolated in this study, two putative new APMV species (*Avian metaavulavirus 24* and *27*) were identified from viruses isolated in this study and from retrospective GenBank sequences, and two putative new APMV species (*Avian metaavulavirus 25* and *26*) were identified solely from retrospective GenBank sequences. Furthermore, co-infections of APMVs were identified in a subset of the samples. The potential limitations of the branch length being the only speciation criterion and the potential benefit of a group pairwise distance analysis are discussed.

**Importance:** Most species of APMVs are understudied and/or underreported and many species were incidentally identified from asymptomatic wild birds; however, the disease significance of APMVs in wild birds is not fully determined. The rapid rise in high-throughput sequencing coupled with avian influenza surveillance programs have identified 12 different APMV species in the last decade and have challenged the resolution of classical serological methods to identify new viral species. Currently, ICTV’s only criterion for *Paramyxoviridae* species classification is the requirement of a branch length >0.03 using a phylogenetic tree constructed from polymerase (L) amino acid sequences. The results from this study identify one new APMV species, propose four additional new APMV species, and highlight that the criterion may have insufficient resolution for APMV species demarcation and that refinement or expansion of this criterion may need to be established for *Paramyxoviridae* speciation.

## Introduction

Paramyxoviruses (family *Paramyxoviridae*) are a diverse group of RNA viruses that infect mammals (including humans), birds, reptiles, and fish. Paramyxoviruses that infect birds belong to the *Avulavirinae* subfamily, whose viruses are referred to as avian paramyxoviruses (APMVs, used hereafter). The International Committee on Taxonomy of Viruses (ICTV) currently defines 22 different avulavirus species that are further divided into three different genera: *Orthoavulavirus* (APMV-1, -9, -12, -13, -16 to -19 [APMV-17, -18, and 19 have also been designated Antarctic penguin viruses A, B, and C, respectively], and -21), *Metaavulavirus* (APMV-2, -5 to -8, -10, -11, -14, -15, -20, and -22), and *Paraavulavirus* (APMV-3 and -4). These species have been reported from a wide range of wild and domestic birds worldwide. The severity of disease (i.e., Newcastle disease) and high mortality rate of virulent APMV-1 (commonly known as Newcastle disease virus) has been responsible for global economic losses in the poultry industry since the 1920s and monitoring programs have detected virulent and avirulent APMV-1 in over 200 avian species (1-3). As a result, APMV-1 is the most studied APMV and has been well characterized with the delineation of two classes (I and II) and at least 22 different genotypes (4). Despite the disease importance of APMV-1 in poultry and widely instituted surveillance efforts, information on the disease ecology and genetic diversity of APMV-2 to -22 is relatively limited. Of note, all but one (APMV-5) APMV species have been isolated from wild birds (5, 6).

Wild birds are natural reservoirs of APMVs (except virulent strains of APMV-1 for which this is not resolved) and the exclusive hosts of many APMVs. The implications for disease in wild birds and ecology costs are understudied as compared to domestic birds due to the apparent low pathogenicity of most APMVs in wild birds (6, 7). While chickens and turkeys infected with APMV-1, -2, -3, -6, and -7 have decreased egg production and mild to severe respiratory disease with variable mortality rates (6, 7), APMVs are often isolated from asymptomatic wild birds sampled during influenza A virus (IAV) surveillance studies (6, 8). Due to an increase in surveillance studies in the early 2000’s, APMV-10 to -22 have been detected across the globe in various wild bird species, including penguins and migratory shorebirds and waterfowl (2, 9-11). Furthermore, the accessibility of next-generation and third-generation sequencing have contributed to a better understanding of the molecular characterization and genetic diversity of APMVs in wild birds (2, 12, 13).

While recent advances in sequencing technology have enhanced the characterization of APMVs, there is still a paucity of public sequence data available for APMV-2 to -22 due to historical studies relying on serotyping methods for identifying species (3, 5, 13). For example, GenBank currently contains less than 160 complete genomes for APMV-2 to -22, whereas there are over 1,200 complete genomes for APMV-1 alone (accessed on 13 Aug 2021). Despite the scarcity of sequences in some species, it is known that the genomes of APMVs are approximately 15–17 kb long and typically comprised of six different open reading frames: nucleocapsid (N), phosphoprotein (P), matrix (M), fusion (F), hemagglutinin-neuraminidase (HN), and polymerase (L). Additionally, the P gene contains an mRNA editing site in which one or two G residues are inserted to produce a V and W protein, respectively (14). By sequencing the HN gene or the F gene, the genetic diversity has been studied in a several non-APMV-1 species: APMV-2 (15, 16), APMV-3 (17), APMV-4 (18-22), and APMV-6 (13, 23, 24), and can be used for predicting antigenicity (13, 25). With the exception of sequencing complete genomes of newly described species, even fewer studies have sequenced the L gene from APMVs, which is currently the gene used by ICTV-recommended methods to speciate paramyxoviruses.

Additionally, while the genetic diversity of some APMV species from wild birds in Europe and Asia has been explored (18, 23, 24), there is limited genetic information on APMVs originating in North America (5, 13, 26-28). Thus, the goal of this study was to use MinION sequencing to detect and genetically characterize APMVs isolated from swabs collected from wild birds during 2016–2018 in the United States.

## Results

Thirty-seven, egg-cultured, influenza-negative, hemagglutination-positive swab samples from blue-winged teal, mallards, and ruddy turnstones were collected in the United States in 2016–2018 (Table 2) and were sequenced with MinION using a random strand-switching method to obtain coding-complete genome sequence of APMV species. Initial BLASTn analysis of the APMV genome consensus sequences from MinION sequencing inferred the presence of APMV-1, APMV-2, APMV-4, APMV-6, and APMV-8 in the samples with nucleotide pairwise identities of 72.74–98.29% to currently available sequences in GenBank. Further analysis using phylogenetic trees constructed with ICTV’s *Paramyxoviridae* methodology using complete L amino acid sequences from the samples and from sequences in GenBank confirmed the presence of APMV-1, APMV-4, APMV-6, APMV-8; however, phylogenetic analysis identified three putative new APMV species among the samples from this study, including re-assigning five APMV-2 and four APMV-6 sequences to these new species. Additionally, two putative new APMV species were identified from sequences already classified in GenBank using ICTV’s *Paramyxoviridae* species demarcation criteria of a branch length (nearest node to tip of branch) of >0.03 or group mean distance >0.06 ± SE (Fig. 1 & S1).

**Fig. 1.**
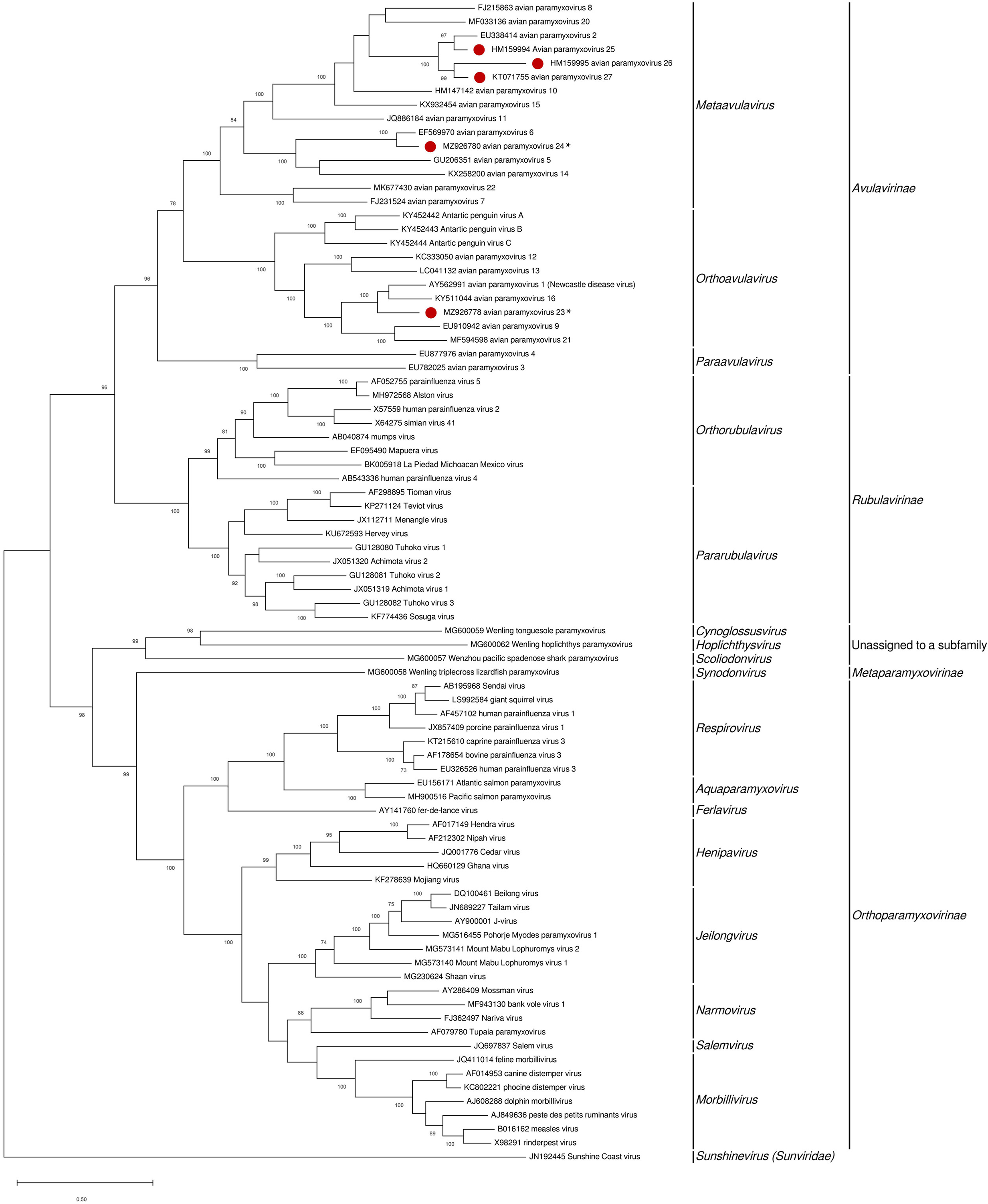
Phylogenetic analysis of complete polymerase (L) amino acid sequences of viruses representing each species in the *Paramyxoviridae* family, including the new species proposed in this study. The evolutionary history was inferred by using the Maximum Likelihood method and JTT matrix-based model (56). The tree with the highest log likelihood (-273062.17) is shown. The percentage of trees in which the associated taxa clustered together is shown next to the branches. Initial tree(s) for the heuristic search were obtained automatically by applying Neighbor-Join and BioNJ algorithms to a matrix of pairwise distances estimated using a JTT model, and then selecting the topology with superior log likelihood value. The tree is drawn to scale, with branch lengths measured in the number of substitutions per site. This analysis involved 84 amino acid sequences. There were a total of 2790 positions in the final dataset. Evolutionary analyses were conducted in MEGA X (55). Red circles denote putative avian paramyxovirus species proposed in this study and asterisks denote isolates sequenced in this study.

**Table 1.**
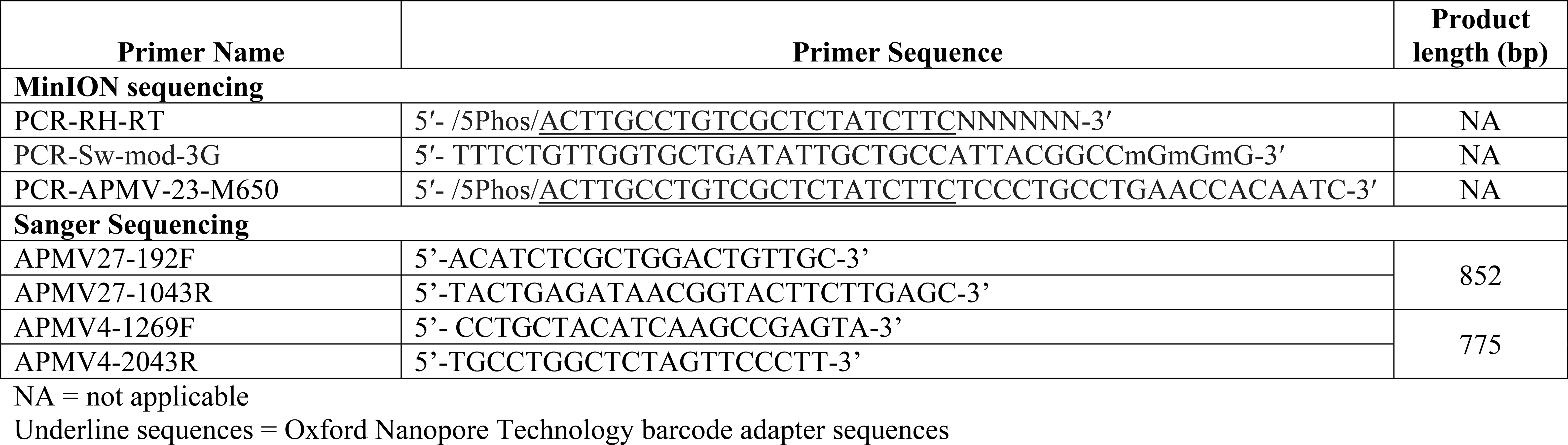
Primers used in this study.

**Table 2.**
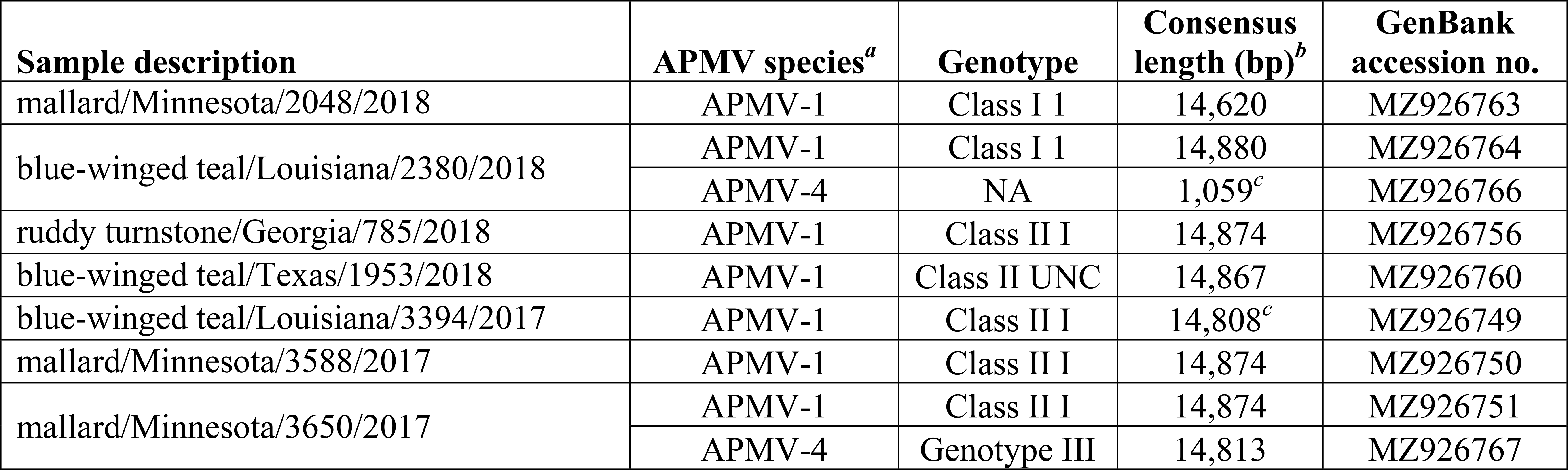
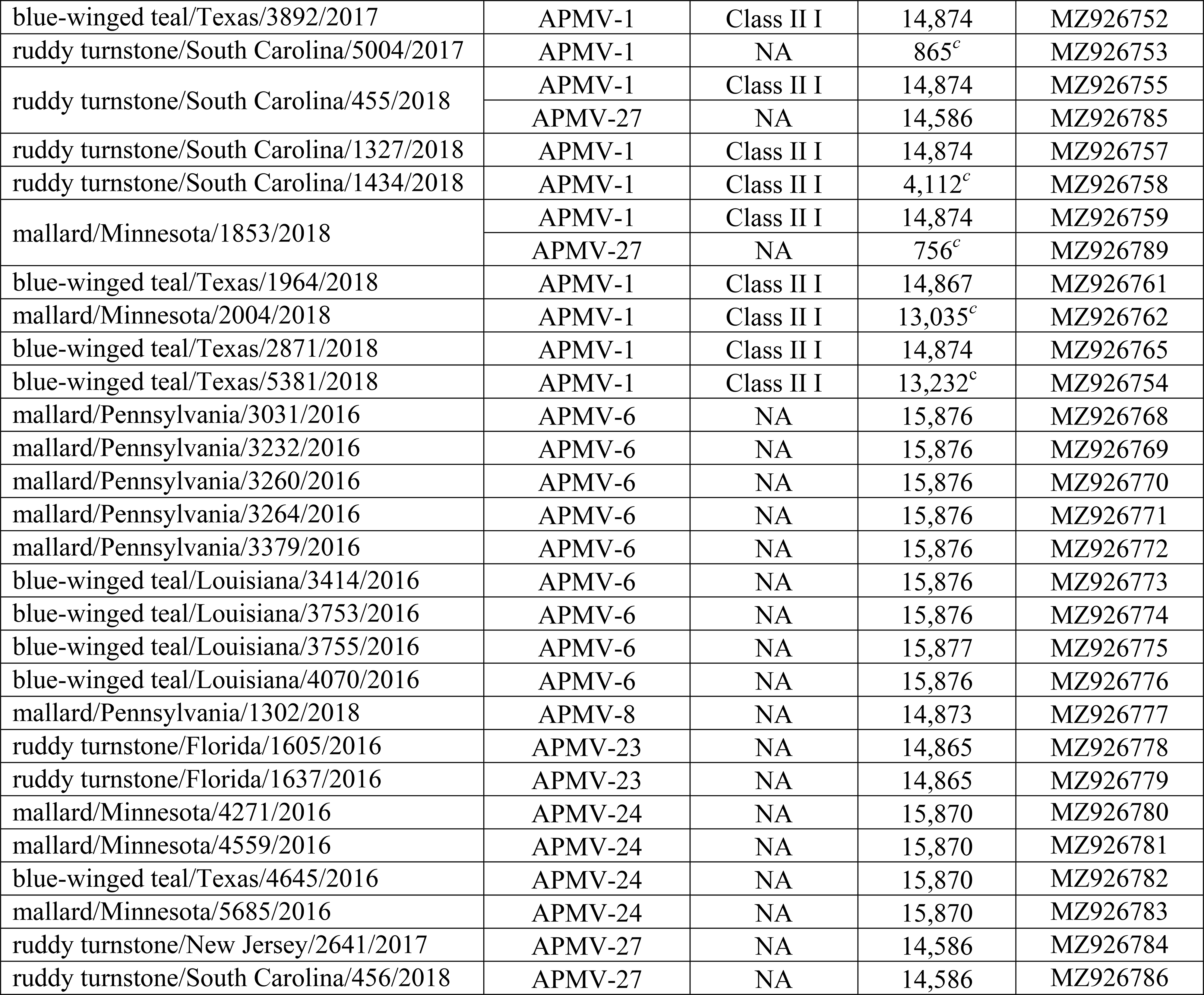
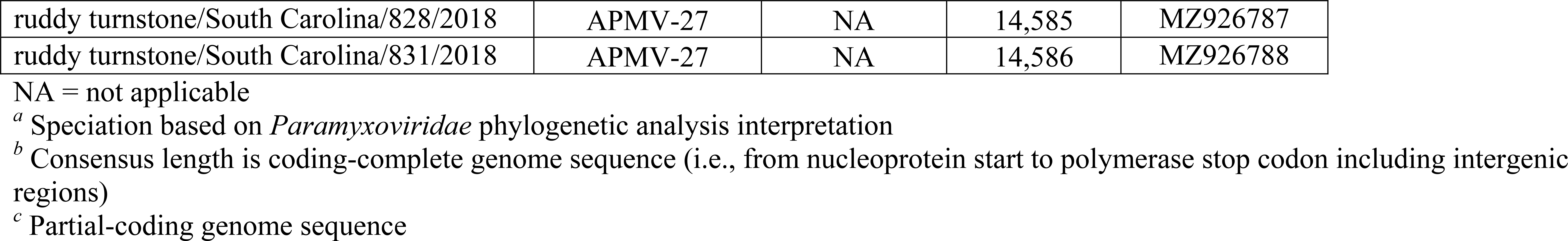
Species and genotypes of avian paramyxoviruses (APMV) sequences isolated from blue-winged teal (*Spatula discors*), mallards (*Anas platyrhynchos*), and ruddy turnstones (*Arenaria interpres*) in the United States between 2016‒2018.

### Putative APMV-23

Using reads from random and targeted strand-switching MinION sequencing, the coding-complete genome sequence of a novel APMV species from two samples collected from ruddy turnstones in Florida (ruddy turnstone/Florida/1605/2016 and ruddy turnstone/Florida/1637/2016) were assembled using a de-novo assembly pipeline, resulting in a 99.96% nucleotide pairwise identity between the two with six nucleotide differences resulting in five amino acid mutations in the NP (T31T, I219L, V221I), F (R105G), and HN (L275F) genes, and one nucleotide change in the NP-P intergenic region. The genomic organization was consistent with most avulaviruses with six transcriptional units (5ʹ-N-P/V/W-M-F-HN-L-3ʹ) of 1,536 nt, 1,188/717/429 nt, 1,095 nt, 1,638 nt, 1,851 nt, and 6,606 nt, respectively. The predicted F protein cleavage site, SGEER↓I, for both samples was monobasic, consistent with typical avirulent APMV strains. GenBank BLASTn analysis of the coding-complete genomes for both samples had the highest nucleotide identity of 72.74% to APMV-1 Goose Paramyxovirus SF02 (GenBank accession: AF473851.2). Phylogenetic analysis of the complete L amino acid sequence using ICTV’s methodology (Fig. 1 & 2) inferred that the novel species is most closely related to APMV-1 and APMV-16 within the *Orthoavulavirus* genus. The novel species formed a major branch with a branch length of 0.151 (Fig. 1 & 2) and according to ICTV’s criterion of a branch length of >0.03 to define a new paramyxovirus species, this branch length supports that these two viruses isolated from ruddy turnstones belong to a novel species, putative *Avian orthoavulavirus-23* (APMV-23).

### APMV-6 and putative APMV-24

During the initial BLASTn data analysis, thirteen isolates had high nucleotide pairwise identities with APMV-6 sequences in GenBank. For nine isolates, collected from blue-winged teal and mallards in Pennsylvania, Texas, and Louisiana, the initial BLASTn analysis showed high nucleotide pairwise identities of 92.39‒99.60% to a single APMV-6 isolated in 2016 from a blue-winged teal in Canada (accession: MW338847.1). However, the remaining four isolates (blue-winged teal/Texas/4645/2016, mallard/Minnesota/4271/2016, mallard/Minnesota/4559/2016, and mallard/Minnesota/5685/2017) (Table 2) had high nucleotide pairwise identities of 95.08–96.01% with other sequences currently designated as APMV-6 in GenBank, but different from the prototypical APMV-6 (GenBank accession: EF569970.1). Because the BLASTn results suggested high diversity between currently assigned APMV-6 and these 13 viruses, another phylogenetic tree was constructed with the newly characterized sample sequences and all 28 APMV-6 sequences currently available in GenBank that had complete L amino acid sequences (Fig. 2). The prototypical APMV-6 cluster contained eleven GenBank APMV-6 sequences (accessions: KP762799.1, MH551526.1, KF267717.1, JX522537.1, KT962980.1, AY029299.1, MW338846.1–7.1, EF569970.1 [APMV-6 reference sequence], EU622637.1, and JN571486.1) which clustered (branch length of 0.065) with the nine sample sequences that best aligned to the 2016 Canadian APMV-6 isolate via BLAST (see above). The phylogenetic analysis showed that the four isolates from this study and 17 of the GenBank sequences currently designated as APMV-6 (accessions: AB759118.1, GQ406232.1, MN163276.1, and MW338848.1–MW338861.1) clustered together and generated a branch length of 0.079 from the prototypical APMV-6 species cluster (Fig. 2). Based on ICTV’s *Paramyxoviridae* speciation criterion (branch length >0.03), these branch lengths suggest that the group of 17 GenBank viruses originally annotated as APMV-6 and the four viruses sequenced in this study constitute a novel species, putative *Avian metaavulavirus-24* (APMV-24).

**Fig. 2.**
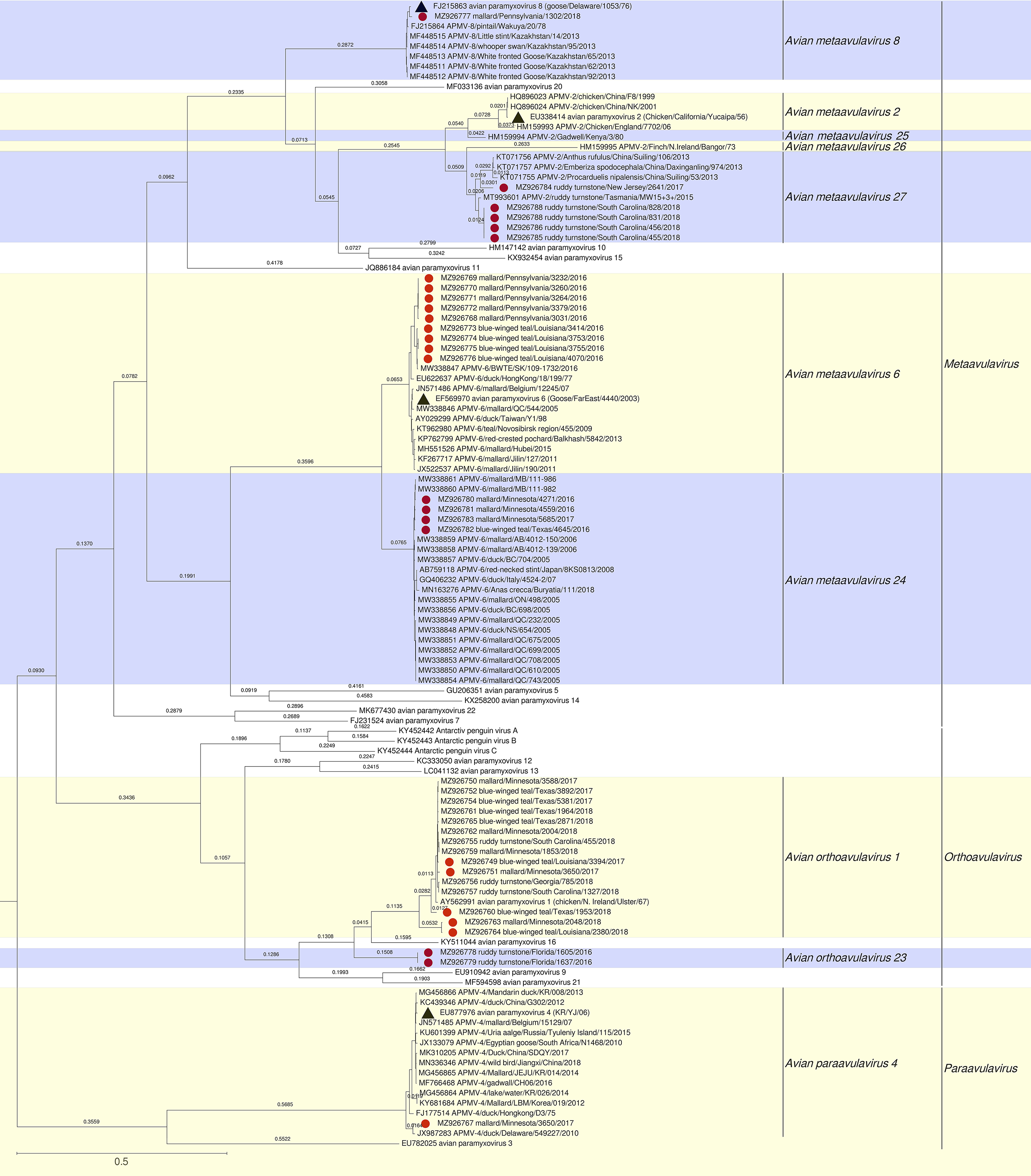
Phylogenetic analysis of all avian paramyxovirus isolates sequenced from this study with viruses in the *Paramyxoviridae* family using complete polymerase (L) amino acid sequences. The evolutionary history was inferred by using the Maximum Likelihood method and JTT matrix-based model (56) with 500 bootstrap replicates. The tree with the highest log likelihood (-281148.93) is shown. Initial tree(s) for the heuristic search were obtained automatically by applying Neighbor-Join and BioNJ algorithms to a matrix of pairwise distances estimated using a JTT model, and then selecting the topology with superior log likelihood value. The tree is drawn to scale, with branch lengths measured in the number of substitutions per site, and branch length values ≥0.010 shown above the branches. This analysis involved 171 amino acid sequences. There were a total of 2773 positions in the final dataset. Evolutionary analyses were conducted in MEGA X (55). Red circles denote samples sequenced in this study and the black triangles denote the GenBank reference sequence for respective avian paramyxovirus species. Avian paramyxovirus species inferred from this study are annotated. For imaging purposes, the full *Paramyxoviridae* phylogenetic tree is not shown (see supplemental Figure 1 and Data Set S4 for full *Paramyxoviridae* phylogenetic analysis).

Coding-complete genome sequence comparison within these two related species showed nucleotide pairwise identities of 93.24‒99.99% for the putative APMV-24 cluster and 85.81‒ 99.98% for the prototypical APMV-6 cluster. The sequence comparison between the prototypical APMV-6 species and the putative APMV-24 species showed low nucleotide pairwise identities of 69.38‒70.40%. The individual amino acid pairwise distances between isolates across the two species were 0.146‒0.154 (Data Set S3) and the between group mean amino acid pairwise distance was 0.150; however, the putative APMV-24 coding-complete genome is organized like a prototypical APMV-6 with seven transcriptional units (5ʹ-N-P/V/W-M-F-SH-HN-L-3ʹ). The N, P, M, SH, HN, and L CDS lengths for the putative APMV-24 sequences were predicted to be the same size as those for the APMV-6 sequences (1,398 nt, 1,293 nt, 1,101 nt, 429 nt, 1,842 nt, and 6,726 nt, respectively). The F CDS lengths differed, with the prototypical APMV-6 species having 1,668 nt and the putative APMV-24 predicted to be 1,683 nt. Additionally, all of the sequences that clustered with the prototypical APMV-6 have predicted F protein cleavage sites of APEPR↓L and all of the putative APMV-24 sequences have IREPR↓L predicted cleavage sites. Other differences include the lengths of the predicted V and W CDSs. The predicted V CDS lengths for all three putative APMV-24 sample sequences from Minnesota and for the GenBank sequences were consistently predicted to be 816 nt; however, the V CDS for the blue-winged teal/Texas/4645/2016 isolate was predicted to be 810 nt. The APMV-6 predicted V CDS lengths from the sample sequences were 807 nt, and the GenBank sequences were 807 or 825 nt. All of the predicted W CDS lengths from the four putative APMV-24 sample sequences were 495 nt, similar to the putative APMV-24 GenBank sequences that had W CDS lengths of 474 or 495 nt. The APMV-6 samples from Pennsylvania uniquely had predicted W CDS lengths of 570 nt, whereas the sample sequences from Texas and Louisiana had predicted W CDS lengths of 489 nt. The GenBank APMV-6 sequences were predicted to have W CDS lengths of 489, 534, and 594 nt.

### APMV-2 and putative APMV-25, -26, and -27 speciation

Five of the studied isolates most closely aligned to an APMV-2 sequence in GenBank during the BLAST-based data analysis, but subsequent phylogenetic analysis showed these isolates were significantly different from the APMV-2 reference sequence (GenBank accession: EU338414). A phylogenetic tree was constructed with all ten GenBank sequences currently denoted as APMV-2 with complete L amino acid sequences and the sample sequences from this study (Fig. 2). The branch lengths for HM159994 and HM159995 supported that these sequences are different species from each other and also from all other APMV-2 GenBank sequences, putative *Avian metaavulavirus-25* (APMV-25) and *Avian metaavulavirus-26* (APMV-26), respectively. On two occasions, asymmetric branch lengths between clusters were observed with one branch length being > 0.03, while the paired branch length was < 0.03 and branch length values near the threshold of the genetic distance cutoff for *Paramyxoviridae* speciation. For these reasons, confident speciation of the remaining sequences could not be ascertained using the single criterion of branch length. Therefore, individual amino acid pairwise and between group mean distances were estimated using MEGA X (Tables 3 & 4).

**Table 3.** Estimates of polymerase amino acid evolutionary distances for individual viruses phylogenetically speciated as avian paramyxovirus (APMV)-2, -25, -26, and -27.

|  |  | Distance or SE<sup>b</sup> |  |  |  |  |  |  |  |  |  |  |  |  |  |  |
| --- | --- | --- | --- | --- | --- | --- | --- | --- | --- | --- | --- | --- | --- | --- | --- | --- |
|  |  | APMV-2 |  |  |  | APMV<br>-25 | APMV<br>-26 | APMV-27 |  |  |  |  |  |  |  |  |
| APMV isolate<sup>a</sup> |  | YUCA | F8 | NK | 7702 | 3 | BANG | 53 | 106 | 974 | MW15 | 2641 | 455 | 456 | 828 | 831 |
| APMV-2 | YUCA |  | 0.001 | 0.001 | 0.005 | 0.008 | 0.016 | 0.011 | 0.011 | 0.011 | 0.010 | 0.010 | 0.011 | 0.011 | 0.011 | 0.011 |
|  | F8 | 0.003 |  | 0.000 | 0.005 | 0.008 | 0.016 | 0.011 | 0.011 | 0.011 | 0.011 | 0.011 | 0.011 | 0.011 | 0.011 | 0.011 |
|  | NK | 0.003 | 0.000 |  | 0.005 | 0.008 | 0.016 | 0.011 | 0.011 | 0.011 | 0.011 | 0.010 | 0.011 | 0.011 | 0.011 | 0.011 |
|  | 7702 | 0.057 | 0.057 | 0.058 |  | 0.008 | 0.016 | 0.011 | 0.011 | 0.011 | 0.011 | 0.011 | 0.011 | 0.011 | 0.011 | 0.011 |
| APMV-25 | 3 | 0.132 | 0.132 | 0.132 | 0.151 |  | 0.014 | 0.009 | 0.009 | 0.009 | 0.009 | 0.010 | 0.009 | 0.009 | 0.009 | 0.009 |
| APMV-26 | BANG | 0.418 | 0.417 | 0.417 | 0.408 | 0.402 |  | 0.012 | 0.012 | 0.012 | 0.012 | 0.012 | 0.012 | 0.012 | 0.012 | 0.012 |
| APMV-27 | 53 | 0.249 | 0.248 | 0.248 | 0.233 | 0.198 | 0.303 |  | 0.002 | 0.002 | 0.005 | 0.005 | 0.005 | 0.005 | 0.005 | 0.005 |
|  | 106 | 0.240 | 0.239 | 0.240 | 0.225 | 0.190 | 0.298 | 0.013 |  | 0.001 | 0.004 | 0.005 | 0.005 | 0.005 | 0.005 | 0.005 |
|  | 974 | 0.241 | 0.241 | 0.241 | 0.226 | 0.191 | 0.301 | 0.015 | 0.004 |  | 0.004 | 0.005 | 0.005 | 0.005 | 0.005 | 0.005 |
|  | MW15 | 0.234 | 0.232 | 0.233 | 0.218 | 0.163 | 0.293 | 0.062 | 0.053 | 0.054 |  | 0.004 | 0.003 | 0.003 | 0.003 | 0.003 |
|  | 2641 | 0.237 | 0.236 | 0.237 | 0.216 | 0.188 | 0.301 | 0.068 | 0.059 | 0.061 | 0.051 |  | 0.005 | 0.005 | 0.005 | 0.005 |
|  | 455 | 0.235 | 0.234 | 0.234 | 0.217 | 0.164 | 0.299 | 0.069 | 0.059 | 0.061 | 0.018 | 0.056 |  | 0.000 | 0.000 | 0.000 |
|  | 456 | 0.235 | 0.234 | 0.234 | 0.217 | 0.164 | 0.299 | 0.069 | 0.059 | 0.061 | 0.018 | 0.056 | 0.000 |  | 0.000 | 0.000 |
|  | 828 | 0.235 | 0.234 | 0.234 | 0.217 | 0.164 | 0.299 | 0.069 | 0.059 | 0.061 | 0.018 | 0.056 | 0.000 | 0.000 |  | 0.000 |
|  | 831 | 0.235 | 0.234 | 0.234 | 0.217 | 0.164 | 0.299 | 0.069 | 0.059 | 0.061 | 0.018 | 0.056 | 0.000 | 0.000 | 0.000 |  |

<sup>a</sup> Abbreviated isolate descriptions were used for the following GenBank accession no. in order from top to bottom and left to right: EU338414.1, HQ896023.1, HQ896024.1, HM159993.1, HM159994.1, HM159995.1, KT071755.1, KT071756.1, KT071757.1, MZ926784.1, MT993601.1, MZ926785.1, MZ926786.1, MZ926787.1, and MZ926788.
<sup>b</sup> Pairwise distances are shown in the bottom left half in roman. Standard errors (SE) are shown upper right half in italics.

**Table 4.**
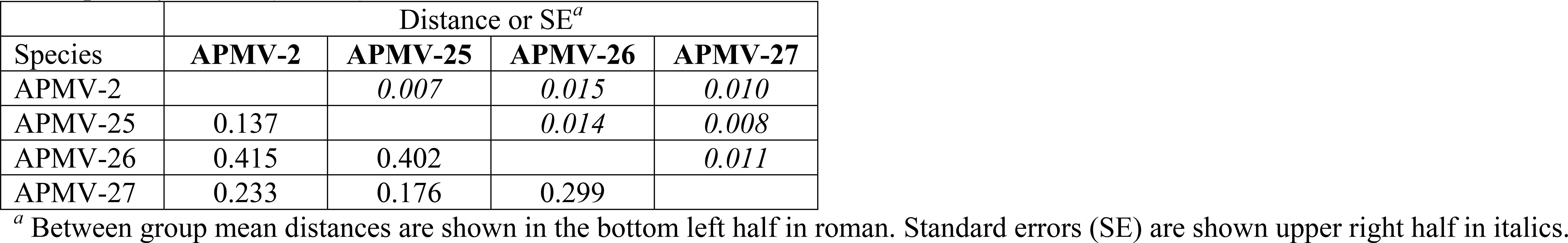
Between group mean estimates of polymerase (L) amino acid evolutionary distances for interspecies comparison for avian paramyxovirus (APMV)-2, -25, -26, and -27.

For the amino acid pairwise and group mean distances a cutoff value has not been declared by ICTV; thus, amino acid pairwise distance values ≥ 0.06 ± SE were interpreted as supporting speciation to account for an average branch length of >0.03. As an example of the asymmetric branch lengths, the branch length from the phylogenetic analysis for HM159993 was 0.037, which suggested speciation from the closest relative, the prototypical APMV-2 cluster that contains the APMV-2 reference sequence. However, the prototypical APMV-2 cluster had a branch length of only 0.02, failing to meet ICTV’s speciation criterion (Fig. 2). The amino acid group mean distances between HM159993 and the prototypical APMV-2 cluster was only 0.058, which suggested that HM159993 is not significantly different to be a different species and was classified as an APMV-2 in this study.

Similarly, four clusters had branch length sums to the closest shared node with some values above or below the 0.03 cutoff: 1) ruddy turnstone/New Jersey/2641/2017 =0.04 (i.e., 0.03 + 0.01), 2) MT993601.1 = 0.01, 3) ruddy turnstone/South Carolina/455/456/828/831/2018 = 0.02, and 4) KT071755.1–7.1 >0.05 (Fig. 2). Additionally, individual amino acid pairwise distances between these nine sequences contained values above and below 0.06 (0.004–0.069, Table 3). After sequentially grouping sequence pairs with the shortest amino acid pairwise distances and re-running the group mean amino acid distances until all groups had > 0.06 distance, the resulting group mean distances demonstrated that these sequences could not be separately speciated from each other, but could be speciated from APMV-2 (group mean distance: 0.23), putative APMV-25 (group mean distance: 0.18), and putative APMV-26 (group mean distance: 0.30) (Table 4); therefore, the five sample sequences and four sequences designated in GenBank as APMV-2 (KT071755-57 and MT993601) represent putative *Avian metaavulavirus 27* (APMV-27).

The complete genome CDS and intergenic regions sequence comparison within the prototypical APMV-2 species cluster showed nucleotide pairwise identities of 94.39‒99.99% and the sequences within the putative APMV-27 species cluster had nucleotide pairwise identities of 82.98‒99.98%. Nucleotide sequence comparison between all APMV-2 and the putative APMV- 25, -26, and -27 showed pairwise identities of 66.58‒87.68%. The coding-complete genome organizations for putative APMV-25, -26, and -27 were similar to APMV-2 with six transcriptional units (5ʹ-N-P/V/W-M-F-HN-L-3ʹ). The P, V, M, and L CDS lengths were identical between all samples within these four species. Within APMV-2, all sequences had identical lengths for all CDSs and the same predicted F cleavage site, KPASR↓F. The predicted F cleavage site and CDS lengths for putative APMV-25 were identical to APMV-2, except for the HN CDS length. Putative APMV-26 varied from APMV-2 and putative APMV-25 in the W, F, and HN CDS lengths, as well as the predicted F cleavage site (LPASR↓F). For the putative APMV-27 sequences, the N, F, and HN CDSs were uniform in length within the species, but differed from APMV-2, putative APMV-25, and -26 except for the HN CDS being the same length with putative APMV-25. The predicted F cleavage sites for putative APMV-27 samples varied within the species (KTAAR↓F, KPTTR↓F, and KPTAR↓F) and were different from that of APMV-2, putative APMV-25 and -26 (Table 5).

**Table 5.** Genome comparisons of avian paramyxovirus (APMV)-2, -25, -26, and -27.

| GenBank<br>Accession | Species | N (nt) | P (nt) | V (nt) | W (nt) | M (nt) | F (nt) | HN (nt) | L (nt) | Predicted F<br>Cleavage Site |
| --- | --- | --- | --- | --- | --- | --- | --- | --- | --- | --- |
| HQ896023.1 | APMV-2 | 1,374 | 1,200 | 699 | 624 | 1,110 | 1,611 | 1,743 | 6,729 | KPASR↓F |
| HQ896024.1 | APMV-2 | 1,374 | 1,200 | 699 | 624 | 1,110 | 1,611 | 1,743 | 6,729 | KPASR↓F |
| EU338414.1 | APMV-2 | 1,374 | 1,200 | 699 | 624 | 1,110 | 1,611 | 1,743 | 6,729 | KPASR↓F |
| HM159993.1 | APMV-2 | 1,374 | 1,200 | 699 | 624 | 1,110 | 1,611 | 1,743 | 6,729 | KPASR↓F |
| HM159994.1 | APMV-25 | 1,374 | 1,200 | 699 | 624 | 1,110 | 1,611 | 1,749 | 6,729 | KPASR↓F |
| HM159995.1 | APMV-26 | 1,374 | 1,200 | 699 | 462 | 1,110 | 1,635 | 1,752 | 6,729 | LPASR↓F |
| KT071755.1 | APMV-27 | 1,368 | 1,200 | 699 | 447 | 1,110 | 1,605 | 1,749 | 6,729 | KTAAR↓F |
| KT071756.1 | APMV-27 | 1,368 | 1,200 | 699 | 447 | 1,110 | 1,605 | 1,749 | 6,729 | KTAAR↓F |
| KT071757.1 | APMV-27 | 1,368 | 1,200 | 699 | 447 | 1,110 | 1,605 | 1,749 | 6,729 | KTAAR↓F |
| MZ926784.1 | APMV-27 | 1,368 | 1,200 | 699 | 462 | 1,110 | 1,605 | 1,749 | 6,729 | KPTTR↓F |
| MT993601.1 | APMV-27 | 1,368 | 1,200 | 699 | 462 | 1,110 | 1,605 | 1,749 | 6,729 | KPTAR↓F |
| MZ926785.1 | APMV-27 | 1,368 | 1,200 | 699 | 462 | 1,110 | 1,605 | 1,749 | 6,729 | KPTAR↓F |
| MZ926786.1 | APMV-27 | 1,368 | 1,200 | 699 | 462 | 1,110 | 1,605 | 1,749 | 6,729 | KPTAR↓F |
| MZ926787.1 | APMV-27 | 1,368 | 1,200 | 699 | 462 | 1,110 | 1,605 | 1,749 | 6,729 | KPTAR↓F |
| MZ926788.1 | APMV-27 | 1,368 | 1,200 | 699 | 462 | 1,110 | 1,605 | 1,749 | 6,729 | KPTAR↓F |

F = fusion; HN = hemagglutinin-neuraminidase; L = polymerase; N = nucleocapsid; M = matrix; and P = phosphoprotein.

### APMV-1

Of the 37 samples sequenced, 17 isolates from all three host species and from multiple states, had pairwise identities of 87.8‒98.5% to APMV-1 sequences in GenBank during the initial BLAST-based analysis. Further phylogenetic analysis supported that these viruses were APMV-1 (Fig. 2). All isolates speciated as APMV-1 were predicted to have dibasic F protein cleavage sites and leucine at position 117, which is typical of APMV-1 strains of low virulence. Isolates with complete F CDS (n=16) were analyzed. Two isolates were identified as class I genotype 1 and 13 isolates were assigned to class II genotype I (Table 2) (Fig. S2 & S3). The genotype for one isolate, blue-winged teal/Texas/1953/2018, could not be determined using the reference-based genotyping method; however, phylogenetic analysis identified the sample to be class II and most closely related to genotypes II and X with a bootstrap value of 87% at the defining node (Fig. S2). It is of interest to note that the *Paramyxoviridae* speciation phylogenetic analysis showed that the class I and class II isolates formed two tips with branch length values of 0.053 and 0.028, respectively (Fig. 2), with a group mean amino acid distance of 0.09 ± 0.04.

### Co-infected samples and APMV-4

A sample collected from a blue-winged teal in Louisiana and a sample from a mallard in Minnesota were co-infected with APMV-1 and APMV-4 species (Table 2). The coding-complete genome sequences for both APMVs were sequenced from the mallard/Minnesota/3650/2017 isolate and was the only coding-complete genome sequence for APMV-4 sequenced in this study. The APMV-4 sample was predicted to be avirulent with a predicted F cleavage site of DIQPR↓F and it was genotyped as genotype III (Fig. S4). For the blue-winged teal/Louisiana/2380/2018 isolates, the coding-complete genome was sequenced for APMV-1 and a partial genome was detected for APMV-4. Primers were designed to confirm the partial L CDS. The Sanger sequencing products resulted in a 677 nt consensus that had 100% nucleotide pairwise identity to the MinION consensus sequence, confirming the accuracy of the partial MinION consensus. The 1,059 nt partial L CDS MinION consensus sequence had 98.0% percent sequence identity to APMV-4/duck/Delaware/549227/2010 (accession JX987283) on the initial BLASTn analysis (phylogenetic analysis not performed due to being a partial sequence). Additionally, the partial MinION sequence had 99.9% nucleotide pairwise identity to the partial L CDS of the mallard/Minnesota/3650/2017 isolate sequenced in this study. The high nucleotide pairwise identities to existing APMV-4 sequences from this study and GenBank support that this sample contains APMV-1 and APMV-4.

### Co-infected samples and APMV-27

Two additional samples had co-infections, one from a mallard in Minnesota and the other from a ruddy turnstone in South Carolina. Both were co-infected with APMV-1 class II genotype I and putative APMV-27. The ruddy turnstone/South Carolina/455/2018 isolate, was discussed above (see APMV-27 and APMV-1 sections). A partial APMV-27 genome was detected from the mallard/Minnesota/1853/2018 sample. Sanger sequencing was performed on a 756 nt fragment to confirm the partial MinION consensus sequence, resulting in 100% sequence identity with the MinION consensus sequence. The 619 nt partial polymerase MinION consensus sequence had 96.8% sequence identity to APMV-2/Emberizaspodocephala/China/Daxing’anling/974/2013 (accession KT071757.1) on the initial BLASTn analysis (phylogenetic analysis not performed due to being a partial sequence). When compared to the putative APMV-27 L CDS sequences from other samples in this study, the sample had 100% nucleotide pairwise identity, supporting the putative APMV-27 co-infection in this sample.

### APMV-8

One sample collected from a mallard in Pennsylvania had a high nucleotide pairwise identity of 97.4% to an APMV-8 sequence in GenBank that was isolated from a goose in Delaware (accession FJ215863.2). The phylogenetic analyses confirmed that the sample clustered within the APMV-8 species (Fig. 2). The predicted F cleavage site of YPQTR↓L was identical to all currently available (6 Aug. 2021) APMV-8 F sequences. The length of the coding-complete genome sequence was identical (14,873 nt) to all other currently reported (6 Aug. 2021) APMV-8 full genomic sequences, and the lengths of all CDSs are identical except for a single nucleotide polymorphism (A→G) that results in a W CDS prediction of 612 nt instead of 519 nt.

## Discussion

The genetic diversity and detection of many APMV species in wild birds are unexplored compared to APMV-1 in domestic birds due to the greater disease and economic impacts of APMV-1 to the poultry industry. In fact, the increase in identification of APMV species and discovery of many novel APMV species were only incidentally found due to global IAV surveillance in wild bird populations. This paucity of publicly available data of complete viral genome sequences limits the ability to study the genetic variability among APMV species and track the evolutionary changes, which is required for the design of species-specific diagnostic tests. Therefore, the purpose of this study was to use MinION sequencing to obtain the complete-coding genome sequences of archived hemagglutination-positive, IAV-negative samples collected in the United States from different species of wild birds. Out of the 37 samples sequenced, 41 different APMV sequences were identified with four samples having co-infections. Thirty-five of these isolates had complete L CDS that were speciated using ICTV’s *Paramyxoviridae* phylogenetic methodology and sequences, which identified APMV-1, -4, -6, - 8, and three putative novel APMV species from the samples. In addition, the complete set of APMV-1, 2, 4, 6, and 8 sequences with complete L CDS available in GenBank were evaluated and additional putative APMV species were identified therein.

The first putative novel APMV species, APMV-23, was detected from two ruddy turnstones located in Nassau County, Florida U.S. The coding-complete genome sequences of these two isolates had high nucleotide pairwise identity, which is consistent with these samples being collected on the same day and location. Speciation using ICTV’s methodology showed the sample sequences were most closely related to APMV-16 and APMV-1 and the branch lengths generated from the phylogenetic analysis were sufficient to genetically differentiate these isolates from existing species. Based on the sequence comparison and phylogenetic analysis with ICTV’s classification criterion, the two isolates sequenced in this study should be considered a new species, *Avian orthoavulavirus 23* (genus *Orthoavulavirus*, subfamily *Avulavirinae)*.

The predicted F cleavage sites for both putative APMV-23 viruses were consistent with typical low virulent APMVs (29). However, because other genomic changes have been shown to increase the virulence of APMV-1 (30-32), viral characterization studies will be needed to determine the true disease potential to chickens and wild bird species. This is especially true in that even less is known about the genetic determinants of pathogenicity in other APMVs in wild birds.

The initial BLASTn analysis of the consensus sequences showed that thirteen APMVs detected from the samples most closely aligned with APMV-6 sequences in GenBank with high nucleotide pairwise identities. Phylogenetic analysis of the sample sequences (n= 4) and sequences with complete L CDS in GenBank (n = 17) created a cluster of sequences that were sufficiently distant to warrant classification as a new species. Additionally, the predicted F cleavage sites and gene nucleotide lengths varied between the APMV-6 and the putative new species. These differences are consistent with studies that have investigated the antigenic and genetic diversity of viruses originally classified as APMV-6s and concluded that the variations were associated with two genotypes, G1 and G2, within the APMV-6 species (13, 23, 24, 33-35). For example, the hemagglutination inhibition (HI) titers against APMV-6 reference anti-sera were lower for the G2 viruses as compared to the G1 viruses (13, 23, 24). Additionally, previous phylogenetic analysis of the complete F gene including 11 of the GenBank “G2” sequences demonstrated a 44.7% nucleotide distance between the G1 and G2 groups (24), which is notably higher than the 10% cutoff that has been used for APMV-1 genotyping (36). Another study using the F gene and whole genome sequences of only APMV6-like sequences demonstrated strong support for the genetic differentiation of the “G1” and “G2” samples. While these genetic and antigenic differences were noted, previous studies did not phylogenetically analyze the L gene in the context of all other avulaviruses. The data presented herein was obtained utilizing the ICTV speciation method and demonstrate that the G1 sequences cluster with the prototypical APMV-6 reference strain in this study, and the G2 sequences are within the putative new species cluster. Considering the data obtained herein using ICTV’s paramyxovirus speciation criteria, it is proposed that the “G2” viruses should be considered a new species, *Avian metaavulavirus 24* (genus *Metaavulavirus*, subfamily *Avulavirinae)*.

Similarly, five APMV sequences from the samples had best BLASTn hits with high nucleotide pairwise identities to APMV-2-annotated sequences in GenBank, but the phylogenetic analysis did not support their classification as APMV-2. In this case, the sample sequences and APMV-2-denoted GenBank sequences with complete L CDSs determined that many sequences generated several branch lengths greater than or near the threshold of ICTV’s paramyxovirus speciation criteria. Individual pairwise and group mean distance estimations using amino acid sequences were conducted to further resolve the genetic diversity and conflicting results of using the branch-length criterion within the APMV-2 cluster. The resulting amino acid pairwise distance values supported three putative novel species (APMV-25, -26, and -27) among the five sample sequences and six GenBank sequences. These results are similar to a previous study that have examined the genetic diversity of the complete genome sequences of APMV-2s and concluded there were potential genotypes among APMV-2s using the Yucaipa (accession EU338414, a.k.a., the APMV-2 reference sequence), England (accession HM159993), Kenya (accession HM159994), and Bangor (accession HM159995) strains (16). Interestingly, the previous study suggested that the Bangor strain is antigenically and genetically distinct to be a different subgroup (16). This difference is also noted in the current study, in which the ICTV’s speciation phylogenetic analysis clustered the Bangor strain on a separate node from the Yucaipa, England, and Kenya strains. Based on the phylogenetic analysis with ICTV’s classification criterion, the viruses previously denoted as APMV-2 in GenBank (HM159994 Gadwall/Kenya/3/80 and HM159995 Finch/N.Ireland/Bangor/73) should be reclassified as new species, *Avian metaavulavirus 25 and 26*, respectively (genus *Metaavulavirus*, subfamily *Avulavirinae*). Additionally, it is proposed that five isolates from this study and four previous APMV-2-denoted GenBank sequences should be classified as an additional new species, *Avian metaavulavirus 27* (genus *Metaavulavirus*, subfamily *Avulavirinae*).

The classification of APMVs within a sample has historically relied on serological methods, primarily HI assays, by using anti-sera of known APMV isolates (13, 25, 37). Although this method has differentiated species that are distantly related, cross-reactivity between some APMV serotypes have been identified and accurate species classification may be problematic due to the rapidly increasing identification of novel paramyxovirus species and the difficulty that creates in terms of laboratories having access to anti-sera for the ever increasing number of APMV species (25). The rise in metagenomic sequencing has shed light on viral diversity and challenges the resolution of historical classification methods to identify new viral species (25, 37, 38). Currently, ICTV’s only criterion for species demarcation within the family *Paramyxoviridae* is based on the topology of a phylogenetic tree of the L protein amino acid sequences with unique species having a branch length value >0.03 (https://talk.ictvonline.org/ictv-reports/ictv_online_report/negative-sense-rna-viruses/w/paramyxoviridae/1194/genus-metaavulavirus). While problems within this paramyxovirus classification method are acknowledged (37), the phylogenetic analysis results from this study highlight two important factors. First, the genetic-based criterion has not been systematically applied to sequences that were speciated prior to the ICTV criterion definition, which was addressed for the historic APMV-2 and APMV-6 GenBank sequences in this study. The second factor is that the branch length criterion alone may be insufficient for defining a new species. For example, the APMV-2s and APMV-2-like clusters had sequences with branch lengths at or near the threshold of the cutoff value and pairs of branches often had one branch length greater than the cutoff and one that was lower. Furthermore, the phylogenetic analysis of APMV-1 sequences from the samples sequenced here generated two clusters, with one tip containing lineage class I genotype 1 isolates and the second containing lineage class II genotype I isolates, with branch length values of 0.532 and 0.0281, which would suggest different species based on the ICTV criterion. The genetic diversity of sequences within APMV-1 has been greatly studied (4) and this diversity complicates the efficient application of a single speciation criterion, which currently would suggest that class I and class II APMV-1 viruses would be considered separated species if there were to only be discovered today. Thus, the results from this study suggest the need for the declaration of more criteria for *Paramyxoviridae* speciation that would account for the inter- and intraspecies variation found within the avulaviruses.

In addition to the several putative novel APMV species identified in this study, there were four samples that were determined to be co-infected with two different APMV species from the MinION based sequencing. One sample collected from a mallard in Minnesota in 2018 and one sample from a ruddy turnstone in South Carolina in 2018 were both discovered to have an APMV-1 and a putative APMV-27. Similarly, a co-infection of APMV-1 and APMV-4 were detected from one sample collected from a blue-winged teal in Louisiana in 2018 and one sample collected from a Mallard in Minnesota in 2017. Co-infections of APMV species in birds is not well studied and this could be due to targeted detection methods (e.g., rRT-PCR) and HI assays being used to classify APMV species during bird health and/or AIV surveillance studies (3, 5, 39, 40). The identification of co-infections of APMV species in wild birds in this study highlights the benefits of using a random sequencing approach method to detect viral species in samples.

Several other known APMV species were also identified in the samples. Of the 37 samples sequenced, 13 isolates were speciated as APMV-1, nine isolates were speciated as APMV-6, and one isolate was speciated as an APMV-8. These results are consistent with other studies that have investigated APMV species in wild birds in the United States and other countries, which have noted that APMV-1, 4, 6, and 8 are commonly detected in wild birds in the Charadriiforme and Anseriforme orders (3, 5, 7, 26). Additionally, the predicted F cleavage sites from the isolates were all monobasic or dibasic, which is consistent with avirulent APMVs and similar to other studies that have analyzed the virulence of APMVs in wild birds (3, 7).

Furthermore, the F CDS sequences that were obtained for APMV-1 isolates were genotyped and identified lineages class I genotype 1 and class II genotype I. One isolate classified as an APMV-1 could not be genotyped using reference-based genotyping method; however, phylogenetic analysis supported that the isolate was a Class II lineage. The isolate generated a distinct branch with high bootstrap support (87%) from genotypes X and II in the phylogenetic analysis, which meets APMV-1 classification criterion #9 of requiring a bootstrap value ≥70% to classify a new genotype or sub-genotype. However, this isolate is only one epidemiological event and does not meet criterion #4 of requiring four or more epidemiological events (4). Additional historic and prospective sequence data may allow a more detailed epidemiological inference and identification of a new genotype. The identification of lineages class I genotype 1 and class II genotype I within the samples is consistent with other studies that have identified genotypes of APMV-1 from wild birds, especially waterfowl (28, 40, 41). The detection and identification of several avirulent APMV sequences further supports that wild birds are reservoirs of low pathogenic APMVs.

In conclusion, the results from this study expand the knowledge of APMV diversity from wild birds, including the identification of putative new APMVs. Additionally, and more importantly, the phylogenetic results of samples in this study demonstrate the need for more robust criteria for *Paramyxoviridae* speciation that accounts for avulavirus diversity and the rapidly increasing use of high-throughput sequencing methods being employed for metagenomic and viral detection studies.

## Materials and Methods

### Sample collection and virus isolation

Archived embryonated chicken egg amnio-allantoic fluid samples were used in this study (42). These samples were collected from a 2016–2018 IAV research and surveillance study by the University of Georgia, Southeastern Cooperative Wildlife Disease Study. Thirty-seven samples (originally from oropharyngeal and cloacal swabs collected from blue-winged teal (*Spatula discors*), mallards (*Anas platyrhynchos*), and ruddy turnstones (*Arenaria interpres*) from eight different states in the United States, Table S1), were selected from the archive based on positive hemagglutination using chicken red blood cells (43) and a negative IAV matrix gene via PCR (44). Hemagglutination-positive and IAV-negative samples were tested for APMVs (45) and APMV-1 (46).

### RNA extraction

Total RNA was extracted from 1 mL of amnio-allantoic fluid with Trizol® LS Reagent (Thermo Fisher Scientific, Waltham, MA, USA) following manufacturer’s protocol. RNA was DNase treated using RNase-Free DNase Set (Qiagen, Hilden, Germany) and purified using RNeasy® MinElute® Cleanup Kit (Qiagen) according to manufacturer’s instructions.

### MinION library preparation and sequencing

RNA extracted from all 37 samples was prepared for MinION sequencing (Oxford Nanopore Technologies (ONT), Oxford, UK) using a randomly primed, strand-switching method as previously described (47). In brief, a strand-switching cDNA synthesis reaction mixture was prepared using the SuperScript™ IV Reverse-Transcriptase kit (Thermo Fisher Scientific) by combining DNase treated RNA (from above), PCR-RH-RT (Table 1) as the reverse transcription primer, and ONT’s strand-switching oligo, PCR-Sw-mod-3G (Table 1). The reactions were incubated and then bead purified using KAPA Pure Beads (KAPA Biosystems, Wilmington, MA, USA). Next, the purified cDNA was PCR amplified and barcoded using ONT’s PCR Barcoding Expansion 1-12 (EXP-PBC001) or PCR Barcoding Expansion 1-96 (EXP-PBC096) kits with LongAmp Taq 2x Mastermix (NEB, Ipswich, MA).

MinION libraries were processed by pooling the barcoded samples by equal volume and following ONT’s 1D amplicon/cDNA by Ligation protocol with the Sequencing Ligation Kit (SQK-LSK109) for dA-tailing and ligating ONT’s sequencing adapter. Final libraries were prepared for sequencing by combining Sequencing Buffer (SQB) and Loading Beads (LB) per ONTs instructions and loaded on FLO-MIN106 R9.4.1 flow cells (ONT) using the MinION Mk1B sequencer (ONT). Sequencing was controlled using MinKNOW v.19.05.0–20.10.3 (ONT) without the live basecalling function.

### MinION sequencing data analysis and consensus building

Real-time basecalling, trimming, and demultiplexing of reads were performed using in-house scripts using the GPU version of Guppy v.3.1.5–4.4.1 (ONT) and Porechop v.0.2.4 (https://github.com/rrwick/Porechop). Raw reads were basecalled using the high accuracy model of Guppy using default parameters with the addition of requiring reads to have a q-score ≥ 7 and allowing calibration strand detection. Then, reads were demultiplexed and sequencing adapters were trimmed with Porechop by requiring reads to have barcodes on each end and 99% adapter identity by aligning 1,000,000 reads to all known adapter sets. Viral reads were taxonomically classified with Centrifuge v.1.0.4 (48) using a custom index that contained complete genome sequences of all possible viruses infecting vertebrates and the chicken genome from NCBI, as of 2 November 2020, using methods previously described (47). Reads were processed and classified by Centrifuge during the sequencing for real-time monitoring of the sequencing run.

For sorting reads to determine APMV species co-infections in each sample, all reads classified as an APMV by Centrifuge were extracted, duplicates were removed, and reads were used for reference-based consensus building using Geneious Prime v.2019.1.3 (Biomatters, Aukland, New Zealand). By using a sequence list of one complete genome sequence per APMV species (10), each cluster of reads from Centrifuge were mapped to the sequence list using the “Map to Reference” tool with medium sensitivity and fine tuning by iterating up to 5 times. The resulting consensus sequences were analyzed with BLASTn by sorting by bit-score and subject coverage. The consensus sequences with the highest bit-score and subject coverage from the BLASTn results were selected as the “top hit”.

Reads from isolates that were identified as APMV-1 or APMV-4 were sorted to determine if multi-genotype coinfections were present using similar methods for speciation. The sequence list for genotyping contained one complete F gene coding sequence (CDS) per main genotype for APMV-4 (20) and for each class I and class II of APMV-1 (4). All reads that aligned to the F CDS sequence from speciation were extracted and then mapped to the genotyping sequence list using the “Map to Reference” tool in Geneious with the same parameters as above. Consensus sequences from each potential genotype were analyzed to identify genotypes using BLASTn with the same criteria as above.

For final reference-based consensus building, the best “top hit” for each isolate from speciation was added to the speciation sequence list and the APMV read clusters from each sample were mapped to the sequence list using the same methods for speciation from above. Final consensus sequences were manually edited for homopolymers (49) that created frameshifts and were required to have ≥ 20× depth of coverage at each base. Using BLASTn, the pairwise identity, bit-score, and subject coverage were evaluated for each consensus sequence to determine the best “top hit”. For consensus sequences with pairwise identities of ≤ 88.0% to sequences in GenBank, the reads were re-mapped once to the initial consensus sequence or used for de-novo assembly. For intra- and interspecies coding-complete genome nucleotide comparisons, sequences were aligned in Geneious with MAFFT v7.450 (50) and pairwise identities were examined.

### De-novo assembly

For the putative novel APMV-23 species, which has no similar sequence in GenBank, the partial (first assembly) and coding-complete (after targeted sequencing, see next section) genome sequences were assembled using a de-novo assembly pipeline. The partial genome was built using the trimmed reads from the randomly primed, strand-switching MinION library. The coding-complete genome sequences were constructed using the trimmed reads from both of the MinION random and targeted strand-switching (described below) libraries. Reads were corrected and assembled using Canu v.1.5 (51) with default parameters with the addition of setting the genome size as 15,000 nt and minimum read length of 700 bp. Contigs were polished with two iterative rounds of Minimap2 v.2.17 (52) with default settings and Racon v.1.4.15 (53) with the following command: racon -m 8 -x -6 -g -8 -w 500 trimmed_reads.fastq minimap_output.paf canu_contig.fasta > racon_output.fasta. Final contigs were further polished using Medaka v.1.0.1 (https://github.com/nanoporetech/medaka) with default parameters using the r941_min_high_g360 model.

### Closing the putative APMV-23 genome CDS

For the putative novel APMV-23 species, in both samples, the randomly primed, strand-switching MinION sequencing obtained partial CDS genomes that were missing approximately 1,000 and 3,000 bases of the 5ʹ end of the genome. A reverse primer targeting the M CDS, PCR-APMV-23-M650, (Table 1) was designed using Geneious (54) and used as the reverse-transcription primer for a cDNA strand-switching reaction as described above with RNA extracted from the samples. Then, the samples were PCR barcoded and MinION sequencing libraries were created as described above.

### Sanger Sequencing and data analysis

Sanger sequencing was performed for two samples that were determined to have co-infections but had low genome coverage for APMV-4 and putative APMV-27. Primers were designed in Geneious using the consensus sequence areas of at least 20× depth of coverage in the L CDS from the MinION libraries (Table 1). Total RNA from each sample was reverse transcribed using SuperScript IV™ First-Strand Synthesis System (Thermo Fisher Scientific) per manufacturer’s protocol with 8 uL of total RNA and random hexamers (50 ng/uL). The cDNA was amplified using DreamTaq PCR Master Mix (2×) (Thermo Fisher Scientific) with 10 uM of each primer (Table 1) following manufacturer’s instructions with the following thermocycling conditions: 95°C for 3 min; 35 cycles of 95°C for 30 sec, 50°C for 30 sec, and 72°C for 1 min; 72°C for 3 min. Amplicons were purified using QIAquick PCR Purification Kit (Qiagen) per manufacturer’s directions and then submitted to GENEWIZ (South Plainfield, NJ, USA) for bidirectional sequencing. Using Geneious, low quality regions (Q≥40) and primers were trimmed from the sequences. Consensus sequences were built from the MinION and Sanger outputs using the MAFFT pairwise alignment tool with default settings in Geneious and analyzed with BLASTn.

### *Paramyxoviridae* speciation phylogenetic analysis

APMV sequences with complete L CDS (n=37, obtained from 35 samples, including two samples with coinfections) from MinION sequencing and sequences from NCBI (22 July 2021) classified as APMV-2 (n=10), APMV-4 (n=14), APMV-6 (n=28), and APMV-8 (n=8) with complete L CDS were used to construct phylogenetic trees in accordance to ICTV’s methodology for *Paramyxoviridae* with ICTV’s sequences (https://talk.ictvonline.org/ictv-reports/ictv_online_report/negative-sense-rna-viruses/w/paramyxoviridae). Sequences were translated in Geneious and then aligned in MEGA X (55) by ClustalW with gap opening penalty of 5 and gap extension penalty of 1 for both pairwise and multiple alignments (Data Sets S1 & S2). Phylogenetic trees were constructed using the Maximum Likelihood model and Jones-Taylor-Thornton (JTT) substitution model (56) with 500 bootstrap replicates in MEGA X. Uniform heterogeneity was used to model evolutionary rates among sites and all gaps and missing data were used. The initial tree was inferred using the Nearest-Neighbor-Interchange (NNI) and BioNJ algorithms.

### Evolutionary amino acid distance estimation

Using the ClustalW alignment of the L amino acid sequences used to build the *Paramyxoviridae* speciation phylogenetic tree (from above), individual pairwise and between group mean pairwise distance substitution estimations were computed using the JTT model with uniform rates among sites and 500 bootstrap replicates in MEGA X. All sites with gaps and missing data were removed from the analysis. While not specified by ICTV, pairwise distances were interpreted as supporting speciation if values were >0.06 ± Standard Error (SE), to account for an average node-to-tip distance of >0.03 (the ICTV speciation metric). For the group mean distance estimation, a series of pairwise distances were performed by sequentially grouping the closest sequences if the distance was ˂0.06 ± SE. The iteration stopped and final groups were determined when all pairwise distances were >0.06 ± SE.

### APMV-1 genotyping phylogenetic analysis

Isolates speciated as APMV-1 with complete F CDS from MinION sequencing were genotyped (Class I n=2; Class II n=14) and used for phylogenetic analyses using the criteria and similar methods as previously described (4) with the Oct_30_2020 data sets from the NDV consortium (https://github.com/NDVconsortium/NDV_Sequence_Datasets). Briefly, sequences were aligned in Geneious using MAFFT v.7.450 (50) with default settings. Then, Maximum-Likelihood trees for Class I and Class II were built using RAxML v.8.2.11 (57) in Geneious using the GTR GAMMA I nucleotide model with the rapid bootstrapping and best-scoring ML tree (-f a) algorithm with 1,000 bootstrap replicates (-N 1000). The parsimony random seed (-p) was set to 456 and the rapid bootstrap random seed number (-x) was set to 123.

### APMV-4 genotyping phylogenetic analysis

For the isolate speciated as APMV-4 with complete F CDS, a phylogenetic tree was constructed using 93 F gene sequences from NCBI (04 April 2021) as previously analyzed (20). Sequences were aligned using ClustalW in MEGA X with default settings. A Maximum Likelihood tree was built using the General Time Reversible model (58) with 1,000 bootstrap replicates in MEGA X. A discrete Gamma distribution was used to model evolutionary rate differences among sites and positions with than less than 95% site coverage were eliminated.

The initial tree was inferred using the NNI and BioNJ methods.

## Data availability

Sequences were deposited in GenBank under accessions MZ926749‒MZ926789 (Table 2).

## Acknowledgments

This project was supported by Agriculture and Food Research Initiative Competitive Grant no. 2018-67015-28306 from the USDA National Institute of Food and Agriculture and funded in part by the National Institute of Allergy and Infectious Diseases, National Institutes of Health, Department of Health and Human Services, under contract HHSN272201400006C. Additional support for this project was supported by Grant no. 5T35OD010433-13 from the Office of Research Infrastructure Programs (ORIP), a component of the National Institutes of Health (NIH) and the contents are solely the responsibility of the authors and do not necessarily represent the official view of ORIP or NIH.

We thank Deborah Carter and Alinde Fojtik (Southeastern Wildlife Disease Study, UGA) for their technical assistance.

The authors declare no potential conflicts of interest with respect to the research, authorship, and/or publication of this article.

## Supplementary material

Supplementary materials for this article is available online.

## Supplementary Tables

**Table S1.** Collection information for each avian paramyxovirus (APMV) isolate sequenced in this study.

| Isolate name<sup>a</sup> | Collection Information |  |  |  |
| --- | --- | --- | --- | --- |
|  | Host | County | State | Collection date (DD-MMM-YYYY) |
| APMV-1/mallard/Minnesota/3588/2017 | *Anas platyrhynchos* | Marshall | Minnesota | 6-Sep-2017 |
| APMV-1/mallard/Minnesota/3650/2017<sup>b</sup> | *Anas platyrhynchos* | Marshall | Minnesota | 8-Sep-2017 |
| APMV-1/blue-winged teal/Texas/3892/2017 | *Spatula discors* | Chambers | Texas | 10-Sep-2017 |
| APMV-1/blue-winged teal/Texas/5381/2017 | *Spatula discors* | Chambers | Texas | 19-Sep-2017 |
| APMV-1/blue-winged teal/Louisiana/3394/2017 | *Spatula discors* | Calcasieu | Louisiana | 23-Sep-2017 |
| APMV-1/ruddy turnstone/South Carolina/5004/2017 | *Arenaria interpres* | Charleston | South Carolina | 10-Dec-2017 |
| APMV-1/ruddy turnstone/South Carolina/455/2018 | *Arenaria interpres* | Charleston | South Carolina | 20-Mar-2018 |
| APMV-1/ruddy turnstone/Georgia/785/2018 | *Arenaria interpres* | Chatham | Georgia | 3-May-2018 |
| APMV-1/ruddy turnstone/South Carolina/1327/2018 | *Arenaria interpres* | N/A | South Carolina | 23-May-2018 |
| APMV-1/ruddy turnstone/South Carolina/1434/2018 | *Arenaria interpres* | Turtle | South Carolina | 23-May-2018 |
| APMV-1/mallard/Minnesota/2004/2018 | *Anas platyrhynchos* | Marshall | Minnesota | 30-Aug-2018 |
| APMV-1/mallard/Minnesota/2048/2018 | *Anas platyrhynchos* | Marshall | Minnesota | 30-Aug-2018 |
| APMV-1/mallard/Minnesota/1853/2018 | *Anas platyrhynchos* | Marshall | Minnesota | 5-Sep-2018 |
| APMV-1/blue-winged teal/Louisiana/2380/2018<sup>b</sup> | *Spatula discors* | La Salle Parish | Louisiana | 22-Sep-2018 |
| APMV-1/blue-winged teal/Texas/2871/2018 | *Spatula discors* | Chambers | Texas | 29-Sep-2018 |
| APMV-4/mallard/Minnesota/3650/2017<sup>b</sup> | *Anas platyrhynchos* | Marshall | Minnesota | 8-Sep-2017 |
| APMV-4/blue-winged teal/Louisiana/2380/2018 | *Spatula discors* | La Salle Parish | Louisiana | 22-Sep-2018 |
| APMV-6/mallard/Pennsylvania/3031/2016 | *Anas platyrhynchos* | Crawford | Pennsylvania | 30-Aug-2016 |
| APMV-6/mallard/Pennsylvania/3232/2016 | *Anas platyrhynchos* | Crawford | Pennsylvania | 9-Sep-2016 |
| APMV-6/mallard/Pennsylvania/3260/2016 | *Anas platyrhynchos* | Crawford | Pennsylvania | 9-Sep-2016 |
| APMV-6/mallard/Pennsylvania/3264/2016 | *Anas platyrhynchos* | Crawford | Pennsylvania | 9-Sep-2016 |
| APMV-6/blue-winged teal/Texas/4645/2016 | *Spatula discors* | Chambers | Texas | 13-Sep-2016 |
| APMV-6/blue-winged teal/Louisiana/4070/2016 | *Spatula discors* | Chambers | Texas | 15-Sep-2016 |
| APMV-6/mallard/Pennsylvania/3379/2016 | *Anas platyrhynchos* | Crawford | Pennsylvania | 15-Sep-2016 |
| APMV-6/blue-winged teal/Louisiana/3414/2016 | *Spatula discors* | La Salle Parish | Louisiana | 19-Sep-2016 |
| APMV-6/blue-winged teal/Louisiana/3753/2016 | *Spatula discors* | La Salle Parish | Louisiana | 23-Sep-2016 |
| APMV-6/blue-winged teal/Louisiana/3755/2016 | *Spatula discors* | La Salle Parish | Louisiana | 23-Sep-2016 |
| APMV-8/mallard/Pennsylvania/1302/2018 | *Anas platyrhynchos* | Mercer | Pennsylvania | 5-Feb-2018 |
| APMV-23/ruddy turnstone/Florida/1605/2016 | *Arenaria interpres* | Nassau | Florida | 7-Feb-2016 |
| APMV-23/ruddy turnstone/Florida/1637/2016 | *Arenaria interpres* | Nassau | Florida | 7-Feb-2016 |
| APMV-24/mallard/Minnesota/4271/2016 | *Anas platyrhynchos* | Roseau | Minnesota | 15-Sep-2016 |
| APMV-24/mallard/Minnesota/4559/2016 | *Anas platyrhynchos* | Marshall | Minnesota | 17-Sep-2016 |
| APMV-24mallard/Minnesota/5685/2017 | *Anas platyrhynchos* | Marshall | Minnesota | 18-Sep-2017 |
| APMV-27/ruddy turnstone/New Jersey/2641/2017 | *Arenaria interpres* | Cape May | New Jersey | 19-May-2017 |
| APMV-27/ruddy turnstone/South Carolina/455/2018<sup>b</sup> | *Arenaria interpres* | Charleston | South Carolina | 20-Mar-2018 |
| APMV-27/ruddy turnstone/South Carolina/456/2018 | *Arenaria interpres* | Charleston | South Carolina | 20-Mar-2018 |
| APMV-27/ruddy turnstone/South Carolina/828/2018 | *Arenaria interpres* | Charleston | South Carolina | 20-Mar-2018 |
| APMV-27/ruddy turnstone/South Carolina/831/2018 | *Arenaria interpres* | Charleston | South Carolina | 20-Mar-2018 |
| APMV-27/mallard/Minnesota/1853/2018<sup>b</sup> | *Anas platyrhynchos* | Marshall | Minnesota | 5-Sep-2018 |

APMV = avian paramyxovirus; N/A = not available.
<sup>a</sup> Speciation based on *Paramyxoviridae* phylogenetic analysis interpretation
<sup>b</sup> Co-infection with different APMV species

## Supplementary Figure Legends

**Fig. S1.** Full phylogenetic analysis of avian paramyxoviruses sequenced from this study with viruses in the *Paramyxoviridae* family using complete polymerase (L) amino acid sequences. The evolutionary history was inferred by using the Maximum Likelihood method and JTT matrix-based model (56) with 500 bootstrap replicates. The tree with the highest log likelihood (-281148.93) is shown. Initial tree(s) for the heuristic search were obtained automatically by applying Neighbor-Join and BioNJ algorithms to a matrix of pairwise distances estimated using a JTT model, and then selecting the topology with superior log likelihood value. The tree is drawn to scale, with branch lengths measured in the number of substitutions per site, and branch length values ≥0.010 shown above the branches. This analysis involved 171 amino acid sequences. There were a total of 2773 positions in the final dataset. Evolutionary analyses were conducted in MEGA X (55). Red circles denote samples sequenced in this study and the black triangles denote the GenBank reference sequence for respective avian paramyxovirus species. Avian paramyxovirus species inferred from this study are annotated.

**Fig. S2.** Full phylogenetic analysis of the complete fusion gene nucleotide sequence of isolate avian paramyxovirus (APMV) 1 class II isolates sequenced in this study. The phylogenetic analysis was constructed using complete fusion gene sequences of 1,901 isolates representing APMV-1 class II genotypes and 14 isolates sequenced in this study. The evolutionary history was inferred by Geneious Prime v.2019.1.3 (Biomatters, Aukland, New Zealand) using the RaxML plugin (57) utilizing the Maximum Likelihood method and GTR GAMMA I model with 1000 bootstrap replicates. The tree is drawn to scale, with branch lengths and measured in the number of substitutions per site. The red circle denotes isolates sequenced in this study.

**Fig. S3.** Full phylogenetic analysis of the complete fusion gene nucleotide sequence of isolate avian paramyxovirus (APMV) 1 class I isolates sequenced in this study. The phylogenetic analysis was constructed using complete fusion gene sequences of 316 isolates representing APMV-1 class I genotypes and two isolates sequenced in this study. The evolutionary history was inferred by Geneious Prime v.2019.1.3 (Biomatters, Aukland, New Zealand) using the RaxML plugin (57) utilizing the Maximum Likelihood method and GTR GAMMA I model with 1000 bootstrap replicates. The tree is drawn to scale, with branch lengths measured in the number of substitutions per site. The red circle denotes isolates sequenced in this study.

**Fig. S4.** Full phylogenetic analysis of complete fusion gene nucleotide sequence of the avian paramyxovirus (APMV) 4 isolate sequenced in this study. The evolutionary history was inferred by using the Maximum Likelihood method and General Time Reversible model (58). The tree with the highest log likelihood (-7279.74) is shown. The percentage of trees in which the associated taxa clustered together is shown next to the branches. Initial tree(s) for the heuristic search were obtained automatically by applying Neighbor-Join and BioNJ algorithms to a matrix of pairwise distances estimated using the Maximum Composite Likelihood (MCL) approach, and then selecting the topology with superior log likelihood value. A discrete Gamma distribution was used to model evolutionary rate differences among sites (5 categories (+G, parameter = 0.8268)). The tree is drawn to scale, with branch lengths measured in the number of substitutions per site. This analysis involved 94 nucleotide sequences. Codon positions included were 1st+2nd+3rd+Noncoding. All positions with less than 95% site coverage were eliminated, i.e., fewer than 5% alignment gaps, missing data, and ambiguous bases were allowed at any position (partial deletion option). There were a total of 1523 positions in the final dataset. Evolutionary analyses were conducted in MEGA X (55). The red circle denotes isolate sequenced in this study.

## Supplementary Data

**Data Set S1.** Alignment file of complete polymerase (L) amino acid sequences used to build the full *Paramyxoviridae* speciation phylogenetic tree with all isolates sequenced in this study.

**Data Set S2**. Alignment file of complete polymerase (L) amino acid sequences used to build the full *Paramyxoviridae* speciation phylogenetic tree with one sequence per species.

**Data Set S3**. Estimates of polymerase amino acid evolutionary distances for individual viruses phylogenetically speciated as avian paramyxovirus (APMV) 6 and 24. Standard errors are shown in italics. Asterisks denote isolates sequenced in this study.

**Data Set S4.** Newick file for Fig. 2 of the full phylogenetic analysis of avian paramyxoviruses sequenced from this study with viruses in the *Paramyxoviridae* family using complete polymerase (L) amino acid sequences.

